# Therapeutic Targeting of Casein Kinase 1δ/ε in an Alzheimer’s Disease Mouse Model

**DOI:** 10.1101/539627

**Authors:** Paula Adler, Janice Mayne, Krystal Walker, Zhibin Ning, Daniel Figeys

## Abstract

Sleep disturbances and memory impairment are common symptoms of Alzheimer’s disease (AD). Given that the circadian clock regulates sleep, hippocampal function, and neurodegeneration, it represents a therapeutic target against AD. Casein kinase 1δ/ε (CK1δ/ε) are clock regulators and overexpressed in AD brains, making them viable targets to improve sleep and cognition. We assessed the effects of a small molecule CK1δ/ε inhibitor (PF-670462) in a cellular model of circadian clocks and in 3xTg-AD mice. Mass spectrometry–based proteomic analyses revealed that PF-670462 treatment *in vitro* upregulated multiple proteins that are downregulated in AD, while administration in 3xTg-AD mice reversed hippocampal proteomic alterations in diverse AD-associated and clock-regulated pathways, including synaptic plasticity and amyloid precursor protein processing. Furthermore, PF-670462 rescued working memory and normalized behavioural circadian rhythms in 3xTg-AD mice. Our study provides proof of concept for CK1δ/ε inhibition and direct clock modulation against AD-related proteomic changes, memory impairment, and circadian disturbances.

## INTRODUCTION

Alzheimer’s disease (AD), the most common type of dementia, is a progressive neurodegenerative disease with cognitive and behavioural symptoms (Livingston et al., 2017). The molecular pathogenesis of AD is complex and several factors have been implicated in AD initiation and progression, including alterations in neurotransmission, amyloid-β (Aβ) levels, cytoskeletal proteins, mitochondrial function, and oxidative stress (Huang and Mucke, 2012; Sanabria-Castro et al., 2017). Given the multifactorial etiology of the disease, it is likely that therapies targeting multiple underlying mechanisms will be most effective. Moreover, disease-modifying approaches may be most beneficial early in the course of AD, before neurodegeneration has progressed and becomes irreversible (Huang and Mucke, 2012).

Memory impairment and disrupted sleep and circadian rhythms are symptomatic hallmarks of AD (Musiek and Holtzman, 2016; Lyketsos et al., 2011). Circadian disturbances, which range from altered sleep timing to severe sleep/wake cycle fragmentation, affect as many as half of people with AD and are a major reason for institutionalization (Lyketsos et al., 2011; Videnovic et al., 2014). Several lines of evidence suggest a bidirectional relationship between circadian dysfunction and AD (Musiek and Holtzman, 2016; Coogan et al., 2013; Videnovic et al., 2014; Chauhan et al., 2017). Disruption of circadian function results from neuropathological changes in the central circadian pacemaker, the suprachiasmatic nucleus (SCN), and may also directly contribute to neurodegeneration in other brain regions through clock-regulated processes such as oxidative stress, proteostasis, neuronal metabolism, and Aβ dynamics (Musiek and Holtzman, 2016). In addition, the circadian system plays critical roles in hippocampal synaptic plasticity and memory (Smarr et al., 2014; Gerstner and Yin, 2010) and circadian disturbances in AD correlate with lower cognitive function, suggesting that clock modulation might represent a novel approach for treating not only sleep problems but also cognitive impairment (Coogan et al., 2013; Bonanni et al., 2005). Given that clock genes throughout the brain have altered expression profiles in AD (Cermakian et al., 2011) and regulate memory, sleep, and neurodegeneration, the circadian clock represents a viable therapeutic target against AD-related memory deficits and sleep disturbances.

Casein kinase 1δ and 1ε (CK1δ/ε) are isoforms of the CK1 family of serine/threonine protein kinases that are implicated in AD and regulation of circadian rhythms (Lee et al., 2009; Perez et al., 2011). Degradation and subcellular localization of core clock proteins are regulated by CK1δ/ε, and mutations in the CK1δ and CK1ε genes in humans and rodents cause behavioural circadian rhythm disorders (Takahashi et al., 2008). Importantly, CK1δ/ε are highly overexpressed in AD brains, indicating that they are therapeutic targets against AD (Perez et al., 2011). A limited number of studies have examined how CK1 inhibition affects Aβ production and tau phosphorylation *in vitro*, and shown that CK1ε overexpression in mouse hippocampus impairs working memory (Perez et al., 2011). However, to our knowledge the global effects of CK1δ/ε inhibition on protein expression have not yet been investigated, and none of these previous reports have addressed whether CK1δ/ε inhibitors can improve cognitive function. Furthermore, whether circadian disturbances in AD can be normalized by inhibiting CK1δ/ε remains unclear.

Here, we assess the therapeutic potential of CK1δ/ε inhibition in an *in vitro* model of circadian clocks and in a triple transgenic mouse model of AD (3xTg-AD mice) with PF-670462, a blood–brain barrier permeable small molecule CK1δ/ε inhibitor that can stabilize circadian rhythms in various mouse models of circadian dysfunction (Badura et al., 2007; Meng et al., 2010). 3xTg-AD mice display memory impairment and circadian abnormalities reminiscent of people with AD, allowing for evaluation of PF-670462 effects (Webster et al., 2014; Sterniczuk et al., 2010). Proteomic analyses using liquid chromatography tandem mass spectrometry (LC-MS/MS) revealed that PF-670462 treatment *in vitro* altered the expression of various AD-related and clock-regulated proteins, while PF-670462 administration *in vivo* rescued hippocampal proteomic alterations in diverse AD-relevant pathways. Moreover, PF-670462 administration restored working memory and normalized behavioural circadian rhythms in 3xTg-AD mice. Our findings provide proof of concept for pharmacological CK1δ/ε inhibition and, more broadly, for circadian clock modulation against AD-related proteomic alterations, cognitive deficits, and circadian disturbances.

## RESULTS

### PF-670462 Treatment Alters Expression of AD-Related and Clock-Regulated Proteins *In Vitro*

To explore the global effects of CK1δ/ε inhibition on protein expression, we first conducted proteomic analysis using LC-MS/MS and Neuro-2a (N2a) mouse neuroblastoma cells, which express multiple clock genes and demonstrate circadian oscillations in gene expression (Chang and Guarente, 2013). We treated N2a cells with either PF-670462 (5 μM) or DMSO (as a control) for 24 h prior to harvesting cells for proteomic analysis. Protein extracts were mixed at a 1:1 ratio with a spike-in standard from heavy-labelled SILAC (stable isotope labelling by amino acids in cell culture) N2a cells to allow for accurate relative quantification (**STAR Methods**). Using a 1% false discovery rate (FDR), a total of 1,595 proteins were identified, of which 1,173 and 1,169 were quantified in cells treated with PF-670462 and DMSO, respectively (**Figure 1A**). Of these, 1,079 proteins yielded relative measurements in a minimum of half of samples (Q50). We used this stringently filtered dataset of accurately quantified proteins for downstream bioinformatic analysis (**Table S1**).

**Figure 1.**
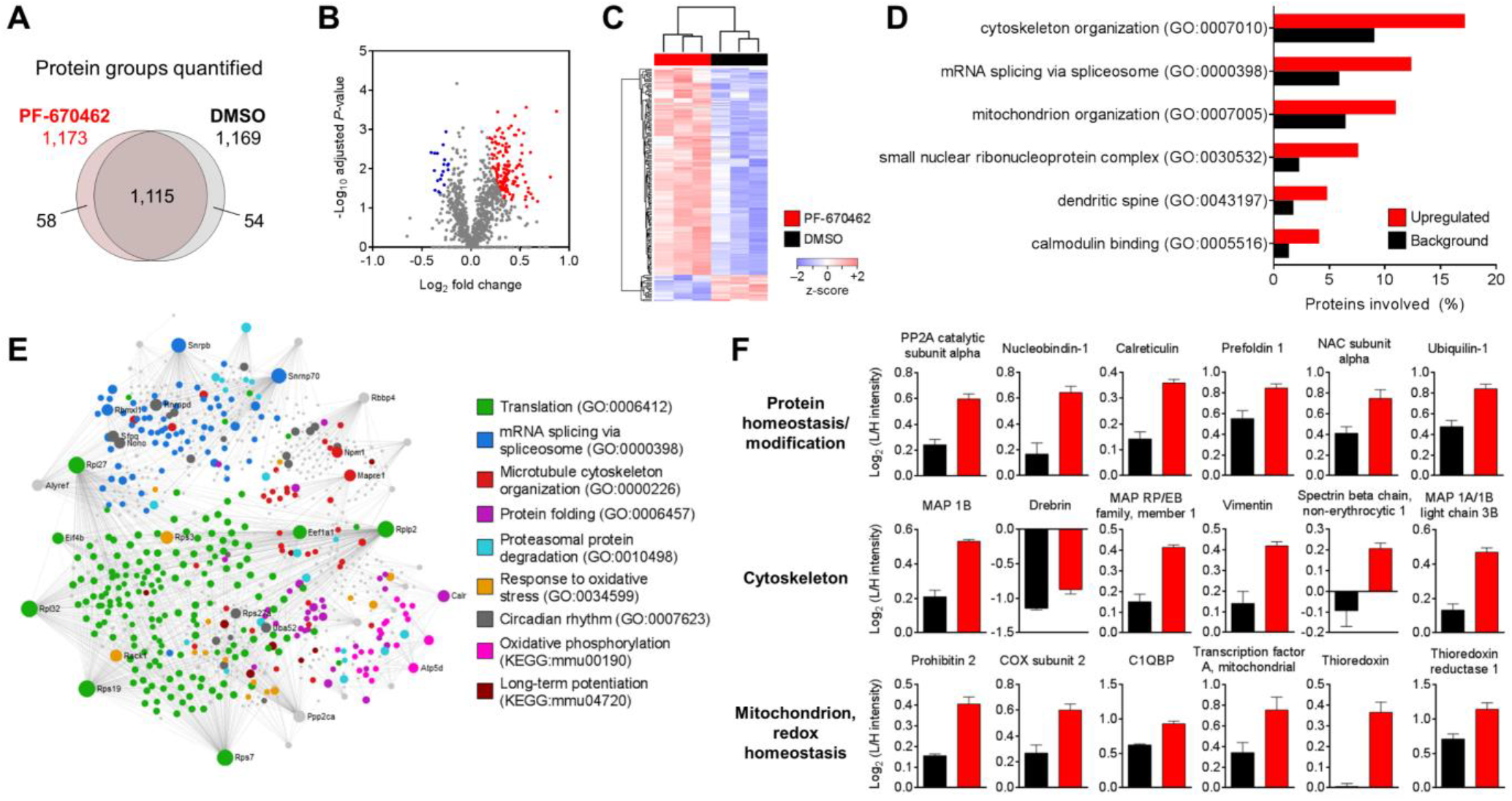
PF-670462 Treatment Alters Expression of AD-Related and Clock-Regulated Proteins *In Vitro*. (A) Proteome coverage: Venn diagram displaying the number of proteins quantified in each treatment condition and overlap between conditions. N2a cells were treated with PF-670462 (5 μM) or DMSO (control) for 24 h and harvested for proteomic analysis. Protein extracts were mixed at a 1:1 ratio with a SILAC spike-in standard from heavy-labelled N2a cells prior to trypsin digestion and LC-MS/MS analysis. n = 3 biological replicates per group. (B) Volcano plot highlighting differentially expressed proteins after PF-670462 treatment (blue = downregulation, red = upregulation; FDR-corrected p < 0.05, two-tailed Student’s *t*-test; s0 = 0.1). (C) Heatmap of z-score normalized abundances of differentially expressed proteins after PF-670462 treatment, showing unsupervised hierarchical clustering of proteins (rows) and samples (columns). (D) GO enrichment analysis using DAVID of proteins upregulated with PF-670462 treatment. A selection of enriched ontology terms associated with AD pathogenesis is shown (p ≤ 0.01, Fisher’s exact test relative to Q50 background). (E) First order PPI network (“subnetwork1”) of proteins differentially expressed after PF-670462 treatment created using the STRING database (interaction confidence score > 0.9). GO and KEGG functional annotations were performed using DAVID and network visualization using NetworkAnalyst. Proteins involved in AD-related and clock-regulated biological processes and pathways shown are represented as coloured nodes. (F) Relative abundances of proteins associated with AD and significantly upregulated in response to PF-670462 treatment. C1QBP, complement C1q binding protein; COX, cytochrome c oxidase; MAP, microtubule-associated protein; NAC, nascent polypeptide-associated complex; PP2A, protein phosphatase 2A. Data are represented as means + SEM. See also Tables S1 and S2.

Comparison of protein abundances (logarithmized L/H ratios) in N2a cells treated with PF-670462 or DMSO showed that the majority (89%, 147 proteins) of differentially expressed proteins were upregulated in response to PF-670462 treatment, with fewer showing downregulation (11%, 19 proteins) (**Figure 1B; Table S2**). Among these differentially expressed proteins were several proteins previously shown to be involved in circadian regulation and to exhibit diurnal rhythms in expression, including casein kinase 2 (Tsuchiya et al., 2009), prohibitin-2 (Kategaya et al., 2012), calreticulin (Noguchi et al., 2017), and galectin-1 (Casiraghi et al., 2010), suggesting that PF-670462 may exert effects on clock-regulated proteins through CK1δ/ε inhibition. To gain insight into the functions of the set of proteins upregulated in response to PF-670462 treatment (top cluster in **Figure 1C**), we performed Gene Ontology (GO) enrichment analysis and identified functional terms that were significantly overrepresented compared to a background of the accurately quantified proteins in our dataset. We found that several of the highly represented functional terms were related to AD pathogenesis, notably cytoskeletal and mitochondrial organization (Huang and Mucke, 2012) (**Figure 1D**).

Given that these biological processes are not only pathologically altered in AD but also known to be regulated by the circadian clock (Chaix et al., 2016; Hoyle et al., 2017), we asked whether other clock-regulated processes and pathways were altered by PF-670462 treatment. Protein– protein interaction (PPI) network analysis of differentially abundant proteins in response to PF-670462 treatment revealed that these proteins take part in a wide range of highly interconnected processes and Kyoto Encyclopedia of Genes and Genomes (KEGG) pathways that have been implicated in AD pathogenesis, including proteasomal protein degradation, response to oxidative stress, long-term potentiation (LTP), oxidative phosphorylation, and circadian rhythm (Manczak et al., 2004; Huang and Mucke, 2012) (**Figure 1E**). Interestingly, these cellular processes also display circadian regulation in mammals (Chaix et al., 2016; Chaudhury et al., 2005), illustrating how the circadian clock and neurodegeneration are closely intertwined and suggesting that clock modulation might protect against neurodegenerative changes at the molecular level. Indeed, we found that PF-670462 treatment resulted in the upregulation of multiple proteins known to be dysfunctional or downregulated in AD and involved in diverse cellular functions, including protein homeostasis and modification, cytoskeletal organization, mitochondrial respiration, and redox homeostasis (**Figure 1F**). Together, these findings suggest that CK1δ/ε inhibition might represent an approach to reverse AD-related protein expression changes across several pathways.

### PF-670462 Administration *In Vivo* Normalizes Hippocampal Proteomic Alterations Associated with AD-Like Pathology

Having established that PF-670462 treatment can alter the expression of AD-related proteins involved in a variety of biological processes *in vitro*, we asked whether PF-670462 administration *in vivo* would have similar effects on the hippocampus, which plays a critical role in memory and displays circadian rhythms in protein expression (Chiang et al., 2017). To examine this, we performed MS-based proteomic analyses using hippocampal tissues from 3xTg-AD mice treated with PF-670462 or vehicle, and non-transgenic (NTg) mice treated with vehicle. Mice were administered daily injections of PF-670462 (30 mg/kg/d) or vehicle for 20 days beginning at 8 months of age, then hippocampal tissues were harvested at two different time points, circadian time (CT) 10 and CT14, on the third day following transfer to constant darkness (DD) (**Figure 2A; STAR Methods**). Samples were processed individually to yield three to five biological replicates per time point for each group, and relative protein abundances were determined by label-free quantification (LFQ). This MS-based analysis identified a total of 1,929 proteins, of which 1,810, 1,608, and 1,551 were quantified in samples from vehicle-treated NTg mice, vehicle-treated 3xTg-AD mice, and PF-670462– treated 3xTg-AD mice, respectively (**Figure 2B**). Of these, ∼1,350 yielded relative measurements in a minimum of half of samples (Q50) at each time point, and we used these filtered datasets for downstream analysis (**Tables S3** and **S4**).

**Figure 2.**
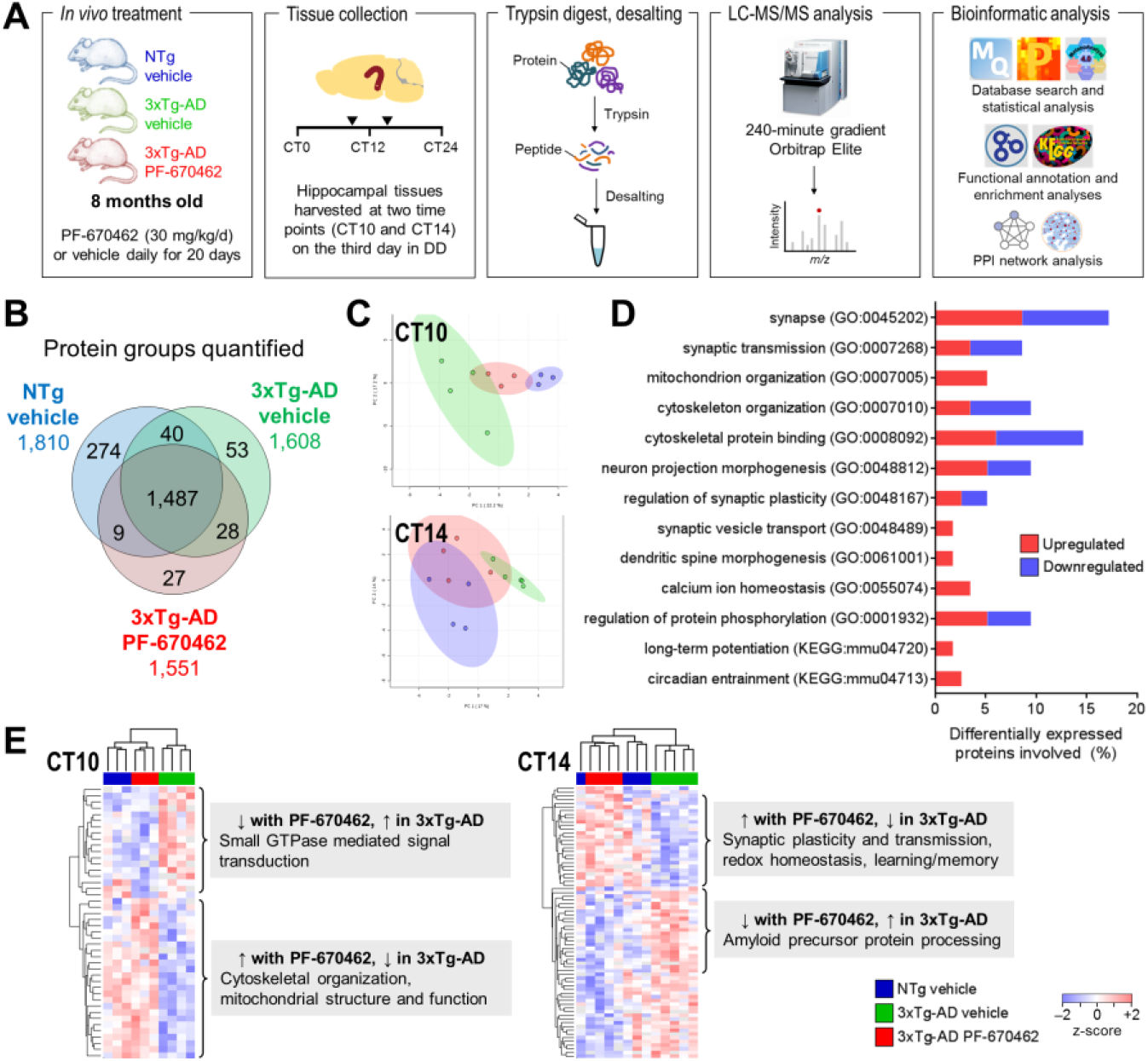
PF-670462 Administration Shifts the Hippocampal Proteomic Profile of 3xTg-AD Mice Towards That of NTg Mice. (A) Study design and workflow: 8-month-old NTg and 3xTg-AD mice were treated daily with PF-670462 (30 mg/kg/d) or vehicle for 20 days, then hippocampal tissues were collected at two time points for proteomic analysis. Protein extracts were digested with trypsin and analyzed by LC-MS/MS. (B) Proteome coverage: Venn diagram displaying the number of proteins quantified in each experimental group and overlap between groups. For CT10: n = 3 NTg vehicle; n = 4 3xTg-AD vehicle; n = 3 3xTg-AD PF-670462. For CT14: n = 4 NTg vehicle; n = 5 3xTg-AD vehicle; n = 4 3xTg-AD PF-670462. (C) PCA of accurately quantified proteins (Q50), showing clustering of samples by experimental group with 95% confidence ellipses. (D) GO and KEGG functional annotations using DAVID of proteins upregulated or downregulated with PF-670462 administration (p < 0.05, two-tailed Student’s t-test; log10 fold change > log101.2 or < –log101.2). A selection of ontology terms and pathways associated with AD pathogenesis is shown. (E) Heatmaps of z-score normalized abundances of differentially expressed proteins after PF-670462 administration, showing unsupervised hierarchical clustering of proteins (rows) and samples (columns). Vehicle-treated NTg and 3xTg-AD mice have distinct hippocampal proteomic profiles, while the profiles of 3xTg-AD mice treated with PF-670462 are similar to those of NTg mice. See also Figure S1 and Tables S3–S6.

On the basis of the expression (logarithmized LFQ intensities) of proteins in these Q50-filtered lists, principal component analysis (PCA) showed that vehicle-treated 3xTg-AD mice can be separated from vehicle-treated NTg mice, while PF-670462 treatment partially reversed proteomic changes associated with AD-like pathology and induced a shift in the hippocampal protein expression profile of 3xTg-AD mice towards that of NTg mice at each time point (**Figure 2C**). Comparison of the 3xTg-AD vehicle-treated and PF-670462–treated groups identified 117 proteins that were differentially expressed in response to PF-670462 treatment (53 upregulated and 64 downregulated; **Tables S5** and **S6**). These proteins take part in a variety of biological processes and pathways, including a number involved in synaptic, cytoskeletal, and mitochondrial functions, as well as circadian entrainment and regulation of protein phosphorylation (**Figure 2D**). Unsupervised hierarchical clustering performed on proteins differentially expressed with PF-670462 administration revealed distinct expression profiles between vehicle-treated 3xTg-AD and NTg mice, reflecting changes associated with AD-like pathology. Moreover, vehicle-treated NTg mice and PF-670462–treated 3xTg-AD mice clustered together, and apart from the vehicle-treated 3xTg-AD group (**Figure 2E**), confirming that widespread alterations in the hippocampal proteome were associated with AD-like pathology and partially normalized by PF-670462 administration.

Interestingly, PF-670462 treatment induced changes in protein expression in distinct molecular pathways at each time point examined, suggesting that circadian regulation might underlie some of the hippocampal responses to CK1δ/ε inhibition. In line with this, we found that differentially expressed proteins were involved in several clock-regulated processes, including cytoskeletal organization and mitochondrial function at CT10, as well as synaptic plasticity and amyloid precursor protein (APP) processing at CT14. PPI network analysis of differentially expressed proteins in the hippocampus of 3xTg-AD mice in response to PF-670462 treatment confirmed the effects of PF-670462 on multiple AD-related processes seen in our *in vitro* study, including cytoskeletal and mitochondrial organization, response to oxidative stress, and proteasomal protein degradation (**Figure 3A**). Administration of PF-670462 in 3xTg-AD mice induced changes in the expression of a greater number of proteins involved in synaptic transmission, regulation of synaptic plasticity, and learning and memory than *in vitro* treatment, which might reflect differences in experimental models at the molecular level as well as contributions of systemic circadian signals *in vivo* (Smarr et al., 2014; Mohawk et al., 2012).

**Figure 3.**
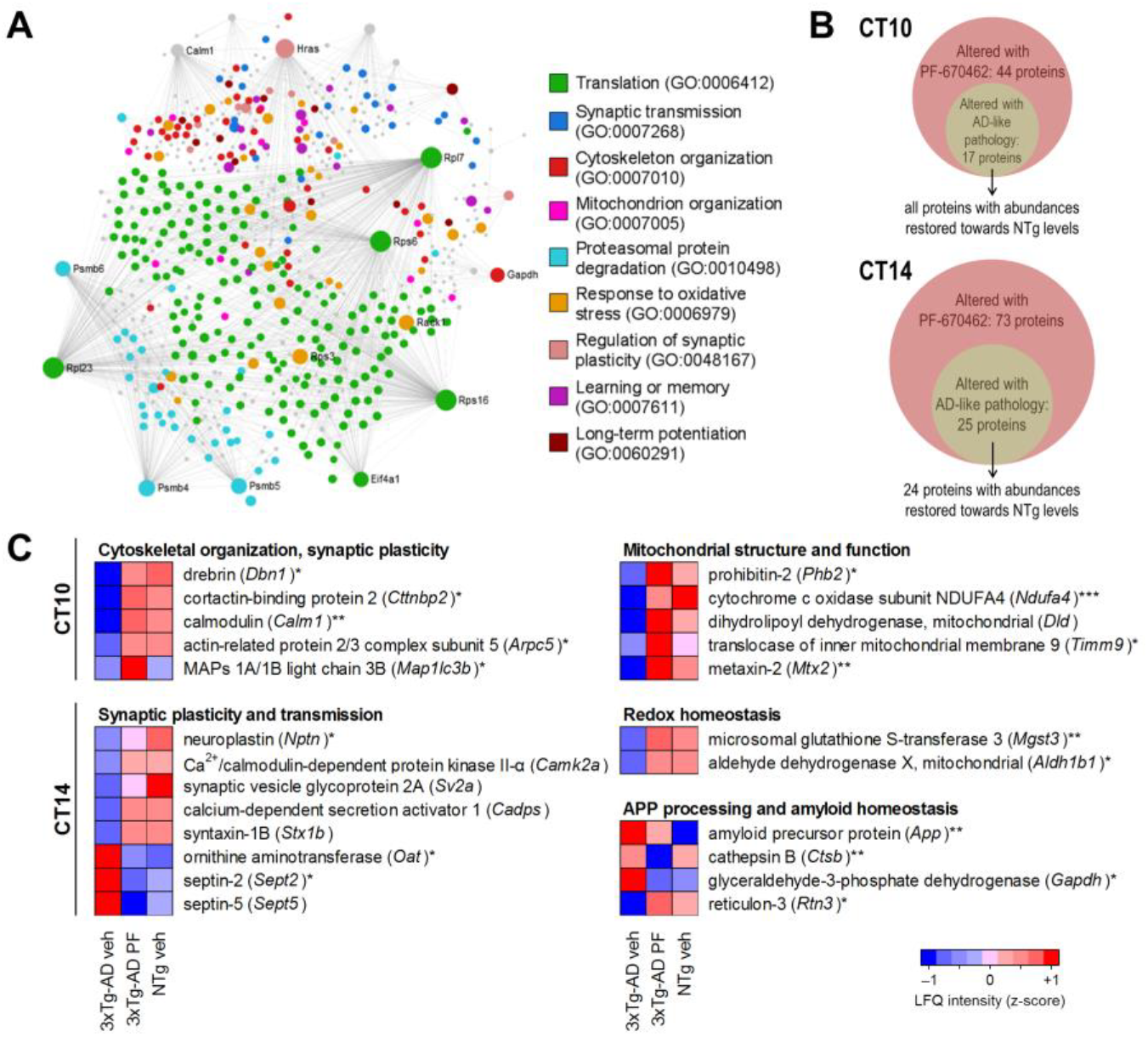
PF-670462 Administration Rescues AD-Related Protein Abundance Changes in the Hippocampus of 3xTg-AD Mice. (A) First order PPI network (“subnetwork1”) of proteins differentially expressed with PF-670462 administration created using the STRING database (interaction confidence score > 0.9) and visualized using NetworkAnalyst (p < 0.05, two-tailed Student’s *t*-test; log10 fold change > log101.2 or < –log101.2). Proteins involved in AD-associated GO biological processes shown are represented as coloured nodes. (B) The number of proteins differentially expressed after PF-670462 administration (3xTg-AD PF-670462 versus 3xTg-AD vehicle), as well as the number of these differentially expressed with AD-like pathology (3xTg-AD vehicle versus NTg vehicle). (C) Relative abundances (mean z-score normalized log10 LFQ intensities) of proteins involved in various AD-related processes. Proteins differentially expressed with PF-670462: *p < 0.05, **p < 0.01, ***p < 0.001, two-tailed Student’s *t*-test; log10 fold change > log101.2 or < –log101.2. MAP, microtubule-associated protein. For CT10: n = 3 NTg vehicle; n = 4 3xTg-AD vehicle; n = 3 3xTg-AD PF-670462. For CT14: n = 4 NTg vehicle; n = 5 3xTg-AD vehicle; n = 4 3xTg-AD PF-670462. See also Figure S1 and Tables S5 and S6.

To gain further insight into the AD-related proteomic alterations in the hippocampus that were reversed by PF-670462 treatment in 3xTg-AD mice, we compared the lists of proteins differentially expressed in vehicle-treated versus PF-670462–treated 3xTg-AD mice, and vehicle-treated 3xTg-AD versus NTg mice at each time point. We found that a substantial proportion of proteins whose abundances were altered with PF-670462 treatment were also differentially expressed with AD-like pathology (39% and 34% at CT10 and CT14, respectively). Strikingly, for all but one of these proteins, PF-670462 treatment normalized their expression levels in the hippocampus of 3xTg-AD mice towards NTg levels (**Figure 3B; Tables S5** and **S6**). To further characterize the effects of PF-670462 on processes implicated in AD pathogenesis, we examined the expression of proteins involved in various functions critical for normal cellular homeostasis and hippocampal function, including cytoskeletal organization, mitochondrial structure and function, synaptic plasticity, and amyloid homeostasis (**Figure 3C**). Administration of PF-670462 in 3xTg-AD mice upregulated the expression of several cytoskeleton-associated proteins, including drebrin and microtubule-associated proteins (MAPs) 1A/1B light chain 3B, as was seen in response to PF-670462 treatment *in vitro*. Hippocampal expression of drebrin, an actin-binding protein that regulates dendritic spine morphogenesis in neurons, is decreased in AD and associated with cognitive function in APP/PS1 mice (Liu et al., 2017). Vehicle-treated 3xTg-AD mice displayed decreased hippocampal levels of drebrin, which were normalized towards NTg levels by PF-670462 treatment. Cortactin-binding protein 2, another actin regulator that plays a role in dendritic arborization and synaptic plasticity (Chen and Hsueh, 2012), was also upregulated in response to PF-670462 treatment in 3xTg-AD mice. Moreover, PF-670462 increased the expression of proteins involved in the hippocampal LTP pathway, including calmodulin and AMPA1 receptor (AMPAR) (**Figure S1A**). PF-670462 administration upregulated the expression of the calcium-buffering protein calmodulin, which was downregulated in the hippocampus of 3xTg-AD mice relative to NTg mice. Decreased expression of calmodulin in AD brains may contribute to deregulated calcium homeostasis, which can lead to neurodegeneration, impairments in synaptic plasticity, and deficits in learning and memory (Marambaud et al., 2009). PF-670462 treatment also upregulated the levels of AMPAR, the trafficking of which is dysregulated by Aβ and represents a critical mechanism underlying LTP induction in the hippocampus (Guntupalli et al., 2016).

Furthermore, PF-670462 administration increased the expression of various proteins involved in mitochondrial function (**Figure 3C**). These included prohibitin-2, which regulates mitophagy and maintenance of mitochondrial structure (Wei et al., 2017; Merkwirth et al., 2012), as well as translocase of inner mitochondrial membrane 9 (Tim9), part of the Tim9-Tim10 complex involved in mitochondrial protein import (Baker et al., 2009). In mice, loss of prohibitin-2 results in impaired mitochondrial respiration and defective mitochondrial ultrastructure, and also leads to widespread neurodegeneration, notably in the hippocampus (Merkwirth et al., 2012). Furthermore, prohibitin-2 is regulated by CK1δ/ε and in turn modulates circadian gene expression (Kategaya et al., 2012), while cortactin-binding protein 2 regulates the actin-binding protein cofilin involved in circadian actin dynamics (Hoyle et al., 2017; Oser et al., 2009), highlighting the effects of PF-670462 on circadian regulation of diverse rhythmic processes. Together, these results suggest that PF-670462 administration *in vivo* normalizes hippocampal expression of multiple proteins linked to AD pathogenesis and implicated in cytoskeletal and mitochondrial functions, consistent with our *in vitro* findings.

We were also interested in examining the effects of PF-670462 treatment on other cellular processes, given that AD-related changes in a variety of processes contribute to neurodegeneration and impaired hippocampal function. In the hippocampus of 3xTg-AD mice, PF-670462 treatment normalized the expression of several proteins implicated in synaptic dysfunction underlying memory impairment in AD, including neuroplastin and septin-2 (**Figure 3C**). Neuroplastin plays an essential role in hippocampal LTP, and neuroplastin deficiency results in impaired associative learning and memory in mice (Bhattacharya et al., 2017). Septin-2, which is increased in AD brains and has been shown to interact with CK1δ (Musunuri et al., 2014; Kategaya et al., 2012), regulates cytoskeletal organization and may take part in the synaptic vesicle cycle (Burre and Volknandt, 2007). In addition to its effects on proteins involved in synaptic plasticity and transmission, PF-670462 upregulated the expression of microsomal glutathione S-transferase 3 (MGST3) and aldehyde dehydrogenase 1B1, which play neuroprotective roles against oxidative stress (Black et al., 2008; Singh et al., 2013) (**Figure 3C**). Interestingly, MGST3 is associated with hippocampal size and is downregulated in AD hippocampus (Ashbrook et al., 2014; Xu et al., 2006). PF-670462 also induced a decrease in the abundance of APP, which has previously been shown to be diurnally regulated (Dobrowolska et al., 2014; Ma et al., 2016) and is thought to play a central role in AD pathogenesis in the amyloid cascade hypothesis (Karran et al., 2011) (**Figure S1B**). Moreover, PF-670462 altered the expression of other proteins involved in APP processing, notably reticulon-3 and cathepsin B. Reticulon-3 was upregulated by PF-670462 and has been shown to inhibit the activity of β-site APP cleaving enzyme (BACE1) (He et al., 2004), which is involved in Aβ production, while cathepsin B was downregulated by PF-670462 and exhibits β-secretase activity (Hook et al., 2008). Inhibitors of BACE1 and cathepsin B have been suggested as therapeutic approaches in AD given their efficacy in reducing Aβ levels and enhancing memory function in preclinical animal models (Hook et al., 2008; Vassar, 2014). Thus, treatment of 3xTg-AD mice with PF-670462 was associated with proteomic changes in hippocampal LTP and APP processing pathways that might be beneficial for AD. Taken together, these results demonstrate that PF-670462 administration *in vivo* can rescue diverse protein expression changes associated with AD in the hippocampus of 3xTg-AD mice, and suggest that CK1δ/ε represent promising therapeutic targets against AD-related hippocampal proteomic alterations.

### PF-670462 Administration Rescues Hippocampal-Dependent Working Memory Deficits in 3xTg-AD Mice

Given that PF-670462 treatment normalized hippocampal proteomic alterations in 3xTg-AD mice, and since CK1ε overexpression has previously been shown to impair working memory in mice (Chen et al., 2017), we hypothesized that PF-670462 administration would improve hippocampal-dependent spatial working memory in 3xTg-AD mice. Moreover, given the associations between anxiety, working memory, and the circadian clock (Wall and Messier, 2000; Lamont et al., 2007), and the influence of anxiety on memory function in AD (Bierman et al., 2009), we were interested in examining the effects of CK1δ/ε inhibition on anxiety in 3xTg-AD mice. To determine whether PF-670462 can reverse AD-associated cognitive deficits, 3xTg-AD and NTg mice were administered daily injections of PF-670462 (20 mg/kg/d) or vehicle starting at 11.5 months of age. Five days after beginning treatment, mice underwent cognitive testing to evaluate working memory and anxiety-like behaviour, with continued daily treatment over the course of testing (**Figure 4A**).

**Figure 4.**
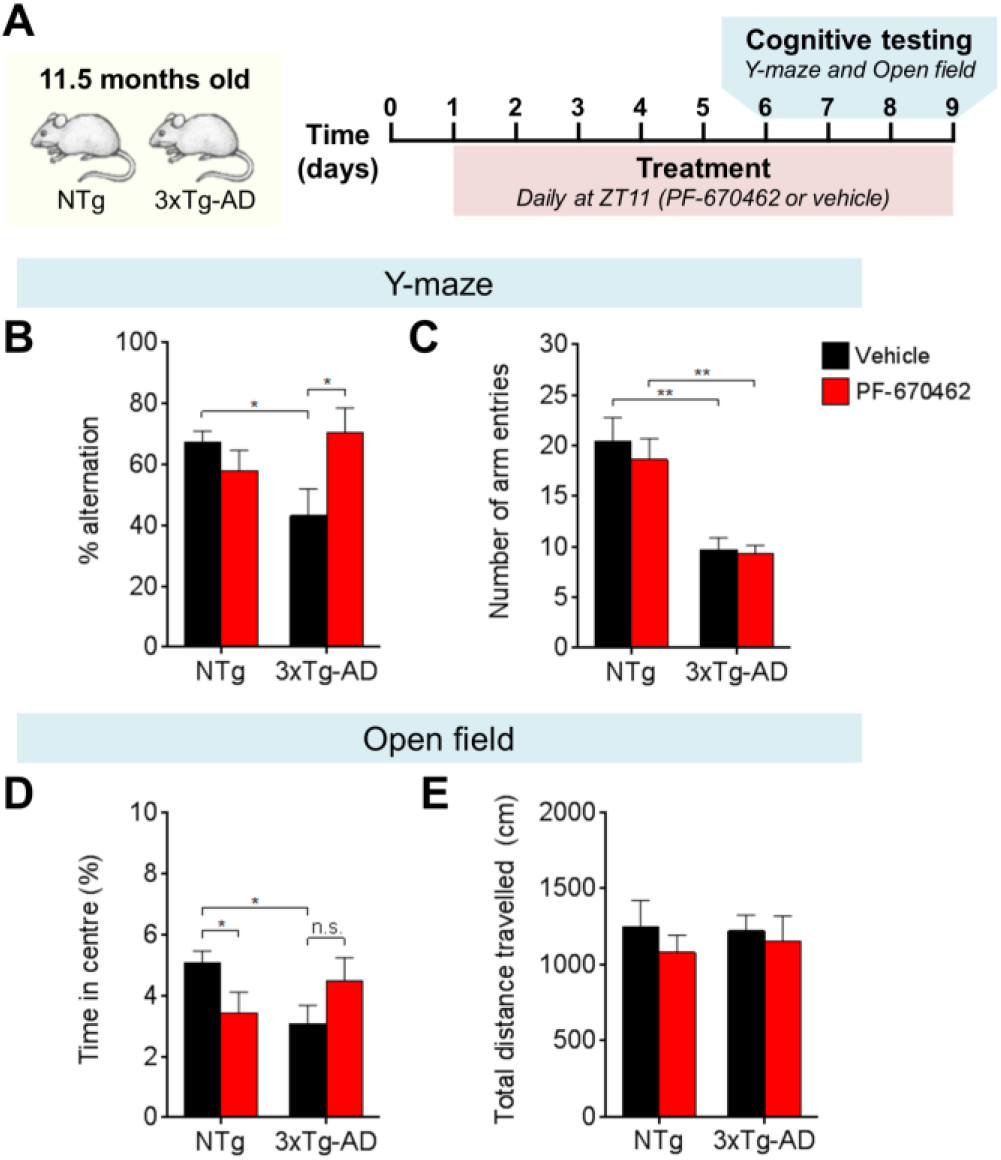
PF-670462 Administration Rescues Working Memory Deficits Without Altering Anxiety-Like Behaviour in 3xTg-AD Mice. (A) Study design: NTg and 3xTg-AD mice were aged to 11.5 months and treatment was started 5 days prior to the beginning of cognitive testing. Mice received daily injections of PF-670462 (20 mg/kg/d) or vehicle, and the Y-maze and open field tests were conducted on days 6 to 9 of treatment. For Y-maze: n = 10 NTg vehicle; n = 11 NTg PF-670462; n = 6 3xTg-AD vehicle; n = 9 3xTg-AD PF-670462. For open field: n = 11 NTg vehicle; n = 11 NTg PF-670462; n = 9 3xTg-AD vehicle; n = 9 3xTg-AD PF-670462. (B) Percent spontaneous alternation during the Y-maze test, showing improvement in spatial working memory in 3xTg-AD mice treated with PF-670462 (*p < 0.05, two-tailed Student’s *t*-test). (C) Number of arm entries during the Y-maze test, showing no changes with PF-670462 treatment (**p < 0.01, two-tailed Student’s *t*-test versus NTg). (D) Percent time spent in the centre during the open field test, showing no change in anxiety-like behaviour in 3xTg-AD mice treated with PF-670462 (*p < 0.05 versus NTg vehicle, two-tailed Student’s *t*-test). (E) Distance travelled during the open field test, showing no differences in locomotion between any groups. Data are represented as means + SEM. n.s., not significant.

To study the impact of PF-670462 treatment on hippocampal-dependent working memory, we assessed spontaneous alternation behaviour in the Y-maze test, a widely used cognitive task to evaluate spatial working memory in rodents. The working memory of 11.5-month-old, vehicle-treated 3xTg-AD mice was significantly impaired compared with age-matched NTg control mice, in line with previous reports (Webster et al., 2014). 3xTg-AD mice treated with PF-670462 showed enhanced working memory relative to vehicle-treated 3xTg-AD mice, while the performance of NTg mice was unaffected by PF-670462 treatment (**Figure 4B**). To confirm that the improvement in working memory seen in 3xTg-AD mice treated with PF-670462 was not due to changes in activity levels, we measured the total number of arm entries and found that it was not altered by PF-670462, with 3xTg-AD mice in both treatment groups showing a similar number of arm entries and fewer than NTg controls (**Figure 4C**). Thus, CK1δ/ε inhibition with PF-670462 rescued hippocampal-dependent working memory deficits in 3xTg-AD mice, and this was associated with normalization of AD-related protein abundance changes in the hippocampus (**Figures 2** and **3**).

To determine whether the enhanced working memory resulting from PF-670462 treatment was associated with changes in anxiety, we also assessed anxiety-like behaviour in the open field test, using the time spent exploring the centre area of the open field as an index of anxiety (Hebda-Bauer et al., 2013). As expected, vehicle-treated 3xTg-AD mice spent less time in the centre compared with NTg control mice, indicating that 3xTg-AD mice exhibit increased anxiety (Webster et al., 2014). Although the time spent in the centre by 3xTg-AD mice treated with PF-670462 was not significantly different from that of vehicle-treated 3xTg-AD mice or NTg mice, PF-670462 treatment seemed to heighten anxiety levels in NTg mice (**Figure 4D**). CK1δ/ε inhibition has previously been shown to alter anxiety-like behaviour in *Clock*Δ19 mice (Arey and McClung, 2012), suggesting that it might act through a similar mechanism to regulate anxiety levels in NTg mice but had no effect on anxiety in 3xTg-AD mice possibly due to pre-existing neurodegeneration in brain regions mediating anxiety. We also measured the total distance travelled during the open field test and did not find significant differences between any groups (**Figure 4E**), indicating that genotype differences in anxiety-like behaviour were unrelated to changes in locomotion. These results suggest that the beneficial effect of PF-670462 on working memory in 3xTg-AD mice was not the result of altered anxiety levels, but is instead linked to treatment-induced changes in the hippocampal proteome. Altogether, our findings provide support for a role of CK1δ/ε in regulating working memory and suggest that CK1δ/ε inhibition could represent a feasible approach to reverse AD-related memory impairment.

### PF-670462 Administration Normalizes Disrupted Behavioural Circadian Rhythms in 3xTg-AD Mice

Previous studies have shown that CK1δ/ε inhibition can restore defective circadian behaviour in various genetic and environmental mouse models of circadian dysfunction (Meng et al., 2010), suggesting that daily treatment with PF-670462 might normalize altered behavioural circadian rhythms in 3xTg-AD mice. 3xTg-AD mice display loss of vasopressin- and vasoactive intestinal polypeptide–expressing neurons in the SCN, resulting in circadian disturbances in patterns of locomotor activity, including a shortened free-running period under DD conditions (Sterniczuk et al., 2010).

To determine whether CK1δ/ε inhibition affects behavioural circadian rhythms in 3xTg-AD mice, we first conducted a preliminary study by administering PF-670462 at 10 and 30 mg/kg/d to allow for assessment of dose– response. At ∼8 months of age, 3xTg-AD mice entrained to a 12-h light:12-h dark (LD) schedule were transferred to DD and their baseline free-running periods determined by monitoring wheel-running activity prior to beginning treatment. Mice were then administered daily injections of PF-670462 (10 or 30 mg/kg/d) or vehicle at CT12, where CT was defined by the zeitgeber time (ZT) of the previous LD schedule (**Figure S2A**). At baseline, 3xTg-AD mice exhibited free-running periods of 23.59 ± 0.1 h as seen on actograms of running wheel activity (**Figure S2B**). Daily treatment with PF-670462, but not vehicle, resulted in lengthening of the free-running period by 0.5 ± 0.26 h and 1.3 ± 0.18 h in 3xTg-AD mice treated with 10 and 30 mg/kg/d PF-670462, respectively (**Figures S2C** and **S2D**). Thus, PF-670462 treatment altered circadian locomotor activity patterns in 3xTg-AD mice, suggesting that CK1δ/ε inhibition might be capable of reversing AD-related circadian disturbances.

Next, we tested whether PF-670462 can re-establish normal behavioural circadian rhythms when administered daily at a moderate dose (20 mg/kg/d) in 3xTg-AD mice. At ∼11 months of age, LD-entrained 3xTg-AD and NTg mice were transferred to DD and their baseline free-running periods determined prior to treatment. Mice were then administered daily injections of PF-670462 (20 mg/kg/d) or vehicle at CT12, as described above (**Figure 5A**). At baseline, 3xTg-AD mice had shortened free-running periods relative to NTg mice (NTg, 23.96 ± 0.02 h; 3xTg-AD, 23.65 ± 0.07 h; **Figures 5B** and **5C**), in line with previous studies (Sterniczuk et al., 2010). Middle-aged NTg mice had free-running periods similar to those seen in aged wild-type mice (Valentinuzzi et al., 1997), resulting in daily activity onsets occurring at approximately the same time each evening, while 3xTg-AD mice displayed irregular activity onsets (**Figure 5D**). Treatment with PF-670462 significantly lengthened the free-running period of NTg mice, as expected (Meng et al., 2010), and also immediately reversed the shortened period in 3xTg-AD mice (**Figures 5E**–**5G**). Furthermore, 3xTg-AD mice treated with PF-670462, but not vehicle, exhibited rapid and stable entrainment of locomotor activity, with regular daily activity onset approximately 18 h after dosing time (**Figures 5B** and **5H**). Thus, daily treatment with PF-670462 normalized disrupted behavioural circadian rhythms associated with AD-like pathology in 3xTg-AD mice. Together, our findings suggest that CK1δ/ε inhibition and, more broadly, direct circadian clock modulation represent viable therapeutic approaches to treat AD-related sleep and circadian disturbances.

**Figure 5.**
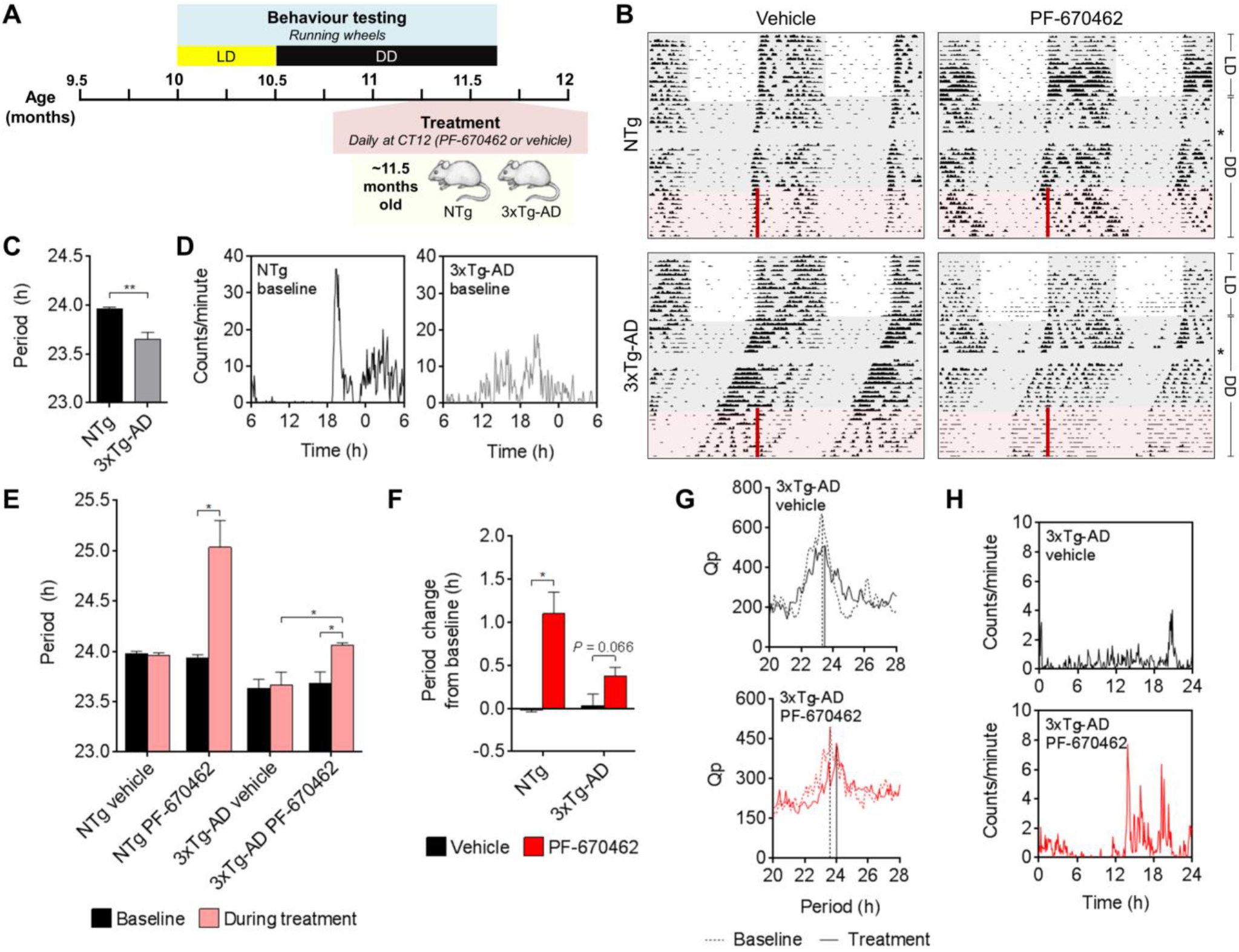
PF-670462 Administration Normalizes Disrupted Behavioural Circadian Rhythms in 3xTg-AD Mice. (A) Study design: Wheel-running activities of NTg and 3xTg-AD mice were recorded for 2 weeks in LD, then mice were released into DD at ∼10.5 months of age. After 3 weeks in DD, mice received daily injections of PF-670462 (20 mg/kg/d) or vehicle for 10 days in DD. n = 5 NTg vehicle; n = 3 NTg PF-670462; n = 6 3xTg-AD vehicle; n = 5 3xTg-AD PF-670462. (B) Representative actograms of wheel-running activities under different lighting conditions of ∼11-month-old NTg and 3xTg-AD mice treated with PF-670462 (20 mg/kg/d) or vehicle daily. The timing of treatment is indicated by red bars. Missing data due to computer failure are indicated by an asterisk. (C) Baseline free-running period under DD of NTg (n = 8) and 3xTg-AD (n = 11) mice (**p < 0.01, two-tailed unpaired Student’s *t*-test with Welch’s correction). (D) Baseline 24-hour activity profiles under DD of NTg and 3xTg-AD mice. Profiles reflect the mean daily activity across 11 days in DD prior to beginning treatment. (E) Free-running period under DD before and during treatment of NTg and 3xTg-AD mice with PF-670462 or vehicle (*p < 0.05 baseline versus during treatment, two-tailed paired Student’s *t*-test; *p < 0.05 vehicle versus PF-670462, two-tailed unpaired Student’s *t*-test with Welch’s correction). (F) Change in free-running period under DD from pre-treatment baseline of NTg and 3xTg-AD mice treated with PF-670462 or vehicle (*p < 0.05, two-tailed unpaired Student’s *t*-test with Welch’s correction). (G) Chi-square periodogram analysis of wheel-running activities of 3xTg-AD mice treated with PF-670462 or vehicle. Representative periodograms of 3xTg-AD mice under DD before and during treatment with vehicle or PF-670462, showing change in period with PF-670462 but not vehicle. (H) 24-hour activity profiles under DD of 3xTg-AD mice treated with vehicle or PF-670462. Profiles reflect the mean daily activity across the entire dosing period. Data are represented as means + SEM. See also Figure S2.

## DISCUSSION

An estimated 44 million people worldwide are living with dementia, the most common cause of which is AD (Collaborators, 2018). There is an urgent need for new therapies that address not only cognitive impairment but also behavioural symptoms commonly seen in AD, including sleep disturbances. However, drug development for AD has proven to be very challenging, evidenced by the fact that there have been no new drugs approved since 2003 for treatment of the disease (Cummings et al., 2018). The multifactorial basis of AD pathogenesis indicates that therapies targeting multiple causal factors will likely be most effective (Huang and Mucke, 2012). Memory, sleep, and neurodegeneration are regulated by the circadian clock (Musiek and Holtzman, 2016; Smarr et al., 2014), suggesting that it is a viable therapeutic target against AD. In light of this, CK1δ/ε inhibition represents a feasible potential disease-modifying approach to treat AD, based on evidence that CK1δ/ε are markedly overexpressed in AD brains, play critical roles in regulating the circadian clock, and might also be involved in tau phosphorylation and controlling Aβ production (Perez et al., 2011; Lee et al., 2009).

In the present study, daily administration of the small molecule CK1δ/ε inhibitor PF-670462 rescued hippocampal-dependent working memory deficits and normalized disrupted behavioural circadian rhythms in middle-aged 3xTg-AD mice. Our findings are consistent with recent work showing that CK1ε overexpression in mouse hippocampus impairs working memory (Chen et al., 2017) and build on previous studies demonstrating that PF-670462 can stabilize circadian rhythms in genetic and environmental models of circadian dysfunction in mice (Meng et al., 2010). Enhanced working memory in 3xTg-AD mice was associated with significant proteomic reprogramming in the hippocampus, suggesting that PF-670462–induced changes in a number of pathways critical for neuronal function and cellular homeostasis resulted in improvements in hippocampal function. Additional studies are needed to explore the effects of CK1δ/ε inhibition on long-term memory, which we were unable to evaluate with the cognitive tests used in the present study.

CK1δ/ε inhibition *in vitro* and *in vivo* altered the expression of proteins involved in diverse processes implicated in AD pathogenesis and hippocampal function, including synaptic plasticity and transmission, redox homeostasis, and APP processing. Notably, administration of PF-670462 in 3xTg-AD mice restored the hippocampal expression of multiple proteins involved in these AD-related processes towards NTg levels, thereby partially rescuing protein abundance changes associated with AD-like pathology. The effects of PF-670462 treatment on the hippocampal proteome are likely in part due to modulation of circadian regulation at the levels of the hippocampal clock as well as central SCN pacemaker (Chauhan et al., 2017). In keeping with this, PF-670462 administration induced changes in distinct clock-regulated processes in the hippocampus of 3xTg-AD mice at different time points. Moreover, the circadian system is known to regulate hippocampal LTP as well as other processes that affect memory and cognitive function, including hippocampal neurogenesis and epigenetic control of gene expression (Gerstner and Yin, 2010; Smarr et al., 2014). Further studies are needed to examine the effects of CK1δ/ε inhibition on circadian regulation of hippocampal neurophysiology in the context of AD, as well as to determine the relative contributions of changes in local hippocampal clock function, systemic circadian control, and clock-independent regulation associated with CK1δ/ε inhibition (Gerstner and Yin, 2010).

There are currently no treatments available that have definitive evidence of effectiveness in treating AD-related sleep disturbances (Livingston et al., 2017). Given that sleep disturbances are a common symptom and major reason for institutionalization in AD (Coogan et al., 2013), there is a critical need for new therapies that act on underlying pathogenic mechanisms to treat these symptoms as well as prevent, halt, or reverse the disease. Multiple lines of evidence suggest that the circadian clock is a feasible therapeutic target to treat sleep disturbances in AD, including the AD-related SCN neurodegeneration linked to circadian and sleep dysfunction (Musiek and Holtzman, 2016) as well as the finding that direct clock modulation can restore defective circadian behaviour in mice (Meng et al., 2010). The effectiveness of PF-670462 in normalizing behavioural circadian rhythm disturbances in middle-aged 3xTg-AD mice provides evidence in support of this hypothesis. Furthermore, administration of PF-670462 has previously been shown to be well-tolerated and produce the same effects on behavioural circadian rhythms in a diurnal non-human primate (Sprouse et al., 2009) as in mice. Thus, this small molecule CK1δ/ε inhibitor represents a promising approach to normalize AD-related sleep and circadian disturbances in humans and could be further optimized to develop more highly potent, selective, and bioavailable drugs for the treatment of AD. Several efforts are already underway to synthesize and optimize different small molecule CK1δ/ε inhibitors for various therapeutic applications and use in basic research, including compounds with enhanced discrimination of δ and ε isoforms (Bibian et al., 2013; Halekotte et al., 2017; Monastyrskyi et al., 2018). Additional preclinical studies are needed to characterize the effects of these newer, more selective and potent CK1δ/ε inhibitors on circadian disturbances, as well as cognitive impairment, in the context of AD.

In summary, treatment of 3xTg-AD mice with the CK1δ/ε inhibitor PF-670462 rescued working memory deficits, normalized behavioural circadian rhythm disturbances, and reversed hippocampal proteomic alterations in diverse AD-related pathways. Collectively, our findings suggest that CK1δ/ε inhibition or, more broadly, direct circadian clock modulation has neuroprotective disease-modifying potential and represents a viable therapeutic avenue to treat cognitive impairment and sleep disturbances in people with AD.

## ACKNOWLEDGEMENTS

We thank Dr. Richard Bergeron for providing 3xTg-AD mice for breeding and the University of Ottawa Behavioural Core for providing testing facilities. D.F. acknowledges a Canada Research Chair in Proteomics and Systems Biology. P.A. acknowledges support from an Alzheimer Society of Canada Biomedical Doctoral Award. The work carried out in this study was supported by the Canadian Institute for Advanced Research and the University of Ottawa.

## AUTHOR CONTRIBUTIONS

P.A. designed the study with input from J.M. and D.F. P.A., K.W., Z.N., and J.M. performed the experiments. P.A. conceived the study, analyzed the data, and drafted the manuscript. D.F., J.M., Z.N., and K.W. reviewed and edited the manuscript. All authors were involved in the interpretation of the data and approved the final version of the manuscript.

## STAR METHODS

### KEY RESOURCES TABLE

| REAGENT or RESOURCE | SOURCE | IDENTIFIER |
| --- | --- | --- |
| Chemicals, Peptides, and Recombinant Proteins |  |  |
| PF-670462 CK1δ/ε inhibitor | Cayman Chemical | Cat#14588 |
| 2-hydroxypropyl-β-cyclodextrin | MilliporeSigma | Cat#H5784 |
| DMEM for SILAC media | AthenaES | Cat#0420 |
| [<sup>13</sup>C<sub>6</sub>, <sup>15</sup>N<sub>2</sub>] L-arginine for SILAC media | AthenaES | Cat#0416 |
| [<sup>13</sup>C<sub>6</sub>, <sup>15</sup>N<sub>2</sub>] L-lysine for SILAC media | AthenaES | Cat#0417 |
| [<sup>13</sup>C<sub>6</sub>, <sup>15</sup>N<sub>2</sub>] L-leucine for SILAC media | AthenaES | Cat#0418 |
| [<sup>13</sup>C<sub>6</sub>, <sup>15</sup>N<sub>2</sub>] L-methionine for SILAC media | AthenaES | Cat#0419 |
| Deposited Data |  |  |
| Proteomics data | PRIDE<br>ProteomeXchange | PXD012281, PXD012282 |
| Experimental Models: Cell Lines |  |  |
| Mouse neuroblastoma: N2a: Neuro-2a | American Type Culture<br>Collection | Cat#CCL-131;<br>RRID:CVCL_0470 |
| Experimental Models: Organisms/Strains |  |  |
| Mouse: NTg: B6129SF2/J | The Jackson Laboratory | Stock#101045;<br>RRID:IMSR_JAX:101045 |
| Mouse: 3xTg-AD: B6;129-Tg(APP<sup>Swe</sup>,tauP301L)1Lfa *Psen1*<sup>tm1Mpm</sup>/Mmjax | The Jackson Laboratory | Stock#34830;<br>RRID:MMRRC_034830-JAX |
| Software and Algorithms |  |  |
| MaxQuant version 1.5.2.8 | Cox and Mann, 2008 | https://maxquant.org/ |
| Perseus version 1.5.5.3 | Tyanova et al., 2016 | https://maxquant.org/perseus/ |
| MetaboAnalyst version 4.0 | Chong et al., 2018 | https://www.metaboanalyst.ca/ |
| DAVID version 6.8 | Huang da et al., 2009 | https://david.ncifcrf.gov/home.jsp |
| NetworkAnalyst | Xia et al., 2015 | https://www.networkanalyst.ca/ |
| Periodogram circadian software | Circadian Rhythm<br>Laboratory | https://www.circadian.org/softwar.html |

## CONTACT FOR REAGENT AND RESOURCE SHARING

Further information and requests for resources and reagents should be directed to and will be fulfilled by the Lead Contact, Daniel Figeys.

## EXPERIMENTAL MODEL AND SUBJECT DETAILS

### Animals

Homozygous 3xTg-AD mice (strain: B6;129-Tg(APPSwe,tauP301L)1Lfa *Psen1tm1Mpm*/Mmjax) (Oddo et al., 2003) were compared to age- and sex-matched non-transgenic (NTg) B6129SF2/J control mice. Mice were obtained from The Jackson Laboratory (Bar Harbor, ME, USA; Stock #34830 for 3xTg-AD and #101045 for NTg; 3xTg-AD mice gift of Dr. Richard Bergeron, University of Ottawa) and maintained in-house by homozygous breeding. Mice were group housed in polycarbonate cages with *ad libitum* access to food and water and maintained on a 12-h light:12-h dark (LD) schedule from weaning. Mice from both genotypes were co-housed unless otherwise indicated. All animal experiments were conducted at the University of Ottawa and approved by the local animal care committee in compliance with institutional and Canadian Council on Animal Care guidelines.

### Stable Isotope Labelling by Amino Acids in Cell Culture

To produce heavy isotope labelled reference proteins, Neuro-2a (N2a) cells (mouse neuroblastoma cell line; American Type Culture Collection [ATCC]; Manassas, VA, USA) were cultured in SILAC (stable isotope labelling by amino acids in cell culture) media at 37°C in a 5% CO_2_ humidified incubator as previously described (Chiang et al., 2017). N2a cells were cultured in customized DMEM (AthenaES; Baltimore, MD, USA) in which the natural arginine and lysine were replaced with heavy [^13^C_6_,^15^N_2_] L-arginine (Arg-10) and [^13^C_6_,^15^N_2_] L-lysine (Lys-8) and supplemented with 10% (v/v) dialyzed fetal bovine serum (FBS) (Gibco; Burlington, ON, Canada), 1 mM sodium pyruvate (Gibco), 28 μg/mL gentamicin (Gibco), and 5 μg/mL Plasmocin prophylactic (InvivoGen; San Diego, CA, USA). Cells were maintained in culture with SILAC media for at least 10 doubling times to allow for complete (>98%) incorporation of the isotopically labelled amino acids into cells.

## METHOD DETAILS

### *In Vivo* CK1δ/ε Inhibition and Tissue Collection

8-month-old female 3xTg-AD and NTg mice group housed by genotype from weaning were administered vehicle (20% [w/v] 2-hydroxypropyl-β-cyclodextrin buffered with 25 mM sodium citrate pH 6.0; MilliporeSigma; Oakville, ON, Canada) or the casein kinase 1δ/ε (CK1δ/ε) inhibitor PF-670462 (30 mg/kg body weight/day; Cayman Chemical; Ann Arbor, MI, USA) subcutaneously (s.c.) daily for 18 days at zeitgeber time (ZT) 11.5 in LD, then transferred to constant darkness (DD) and treated in DD for another two days at circadian time (CT) 11.5, where CT was defined by the ZT of the previous LD schedule. After two days in constant darkness, mice were sacrificed by cervical dislocation under dim red light at CT10 and CT14 on the third day of DD (Chiang et al., 2017). The hippocampi were quickly excised, immediately flash frozen in liquid nitrogen, and stored at –80°C until further processing.

### Proteomic Analysis of Hippocampal Tissues

Protein extracts from hippocampal tissues of individual mice were obtained by mechanical homogenization in lysis buffer containing 4% (w/v) sodium dodecyl sulphate (SDS) in 50 mM ammonium bicarbonate (ABC; pH 8.2) supplemented with complete protease and phosphatase inhibitor cocktails (Roche; Mississauga, ON, Canada), followed by sonication (three 10 s pulses with 30 s on ice between each pulse). Proteins from the resulting supernatant were precipitated in 50% acetone/50% ethanol (EtOH)/0.1% acetic acid at a ratio of 1:5 volumes at –20°C overnight. Proteins were pelleted by centrifugation at 16,000 × *g* for 20 min at 4°C and pellets were washed three times with ice cold acetone. Proteins were resuspended in 8 M urea in 50 mM ABC (pH 8.2). Protein concentrations were determined using the DC Protein Assay (Bio-Rad; Mississauga, ON, Canada). Proteins were reduced by incubating samples with 5 mM dithiothreitol (DTT; MilliporeSigma) for 30 min at 56°C with agitation (245 rpm) and subsequently alkylated with 10 mM iodoacetamide (IAA; MilliporeSigma) for 30 min in darkness at room temperature. Protein digestion was performed by incubation with 40:1 (w/w, protein:enzyme) trypsin (Worthington Biochemical Corporation; Lakewood, NJ, USA) overnight at 37°C with agitation (245 rpm). Samples were acidified using 10% (v/v) trifluoroacetic acid (TFA; MilliporeSigma), then desalted using in-house made C_18_ desalting cartridges (C18 beads: ReproSil-Pur C18-AQ, 10 μm; Dr. Maisch GmbH; Beim Brückle, Germany) and desiccated using a SpeedVac. Peptides were resuspended in 0.1% (v/v) formic acid (FA) for liquid chromatography tandem mass spectrometry (LC-MS/MS) analysis.

### *In Vitro* CK1δ/ε Inhibition

Light N2a cells (ATCC; Manassas, VA, USA) were grown in DMEM (Gibco) supplemented with 10% (v/v) FBS (Gibco), 1 mM sodium pyruvate (Gibco), 28 μg/mL gentamicin (Gibco), and 5 μg/mL Plasmocin prophylactic (InvivoGen) at 37°C in a 5% CO_2_ humidified incubator. For CK1δ/ε inhibition, immediately following a media change for synchronization (Yeom et al., 2010; Nakamura et al., 2016), light N2a cells were treated with PF-670462 (5 μM) or dimethyl sulfoxide (DMSO) as a control for 24 h prior to being harvested for proteomic analysis. Heavy N2a cells were treated for 6 or 24 h with PF-670462 (5 μM) or DMSO prior to being harvested for use as a SILAC spike-in standard.

### SILAC-Based Proteomic Analysis of N2a Cells

Protein extracts from light and heavy N2a cells were obtained by homogenization in lysis buffer containing 8 M urea in 50 mM ABC (pH 8.2) supplemented with complete protease and phosphatase inhibitor cocktails (Roche), followed by sonication (three 10 s pulses with 30 s on ice between each pulse). Protein concentrations were determined using the Bradford assay (Bio-Rad). Lysates from light and heavy N2a cells were mixed at a 1:1 ratio and loaded onto 30-kDa molecular weight cutoff Microcon filters (Millipore; Billerica, MA, USA). Proteins were reduced by incubating samples with 20 mM DTT for 30 min at 37°C with agitation (245 rpm) and subsequently alkylated with 20 mM IAA for 30 min in darkness at room temperature. Protein digestion was performed by incubation with 40:1 (w/w, protein:enzyme) trypsin (Worthington Biochemical Corporation) overnight at 37°C with agitation (245 rpm). Samples were acidified using 10% (v/v) FA, then desalted using in-house made C_18_ desalting cartridges (C18 beads: Dr. Maisch GmbH) and desiccated using a SpeedVac prior to being resuspended in 0.1% (v/v) FA for LC-MS/MS analysis.

### LC-MS/MS Analysis

4 μL of resuspended peptides (equivalent to 2 μg of proteins) from each sample were analyzed by an online reverse-phase LC-MS/MS platform consisting of an Eksigent NanoLC 425 system (AB SCIEX) coupled with an Orbitrap Elite mass spectrometer (Thermo Fisher Scientific; San Jose, CA, USA) *via* a nano-electrospray source. Prior to MS analysis, peptide mixtures were separated by reverse-phase chromatography using an in-house packed ReproSil-Pur C18-AQ column (75 μm internal diameter × 15 cm, 1.9 μm, 200 Å pore size; Dr. Maisch GmbH; Beim Brückle, Germany) over a 240-min gradient of 5–30% buffer B (acetonitrile [ACN] with 0.1% [v/v] FA) at a flow rate of 300 nL/min. The Orbitrap Elite instrument was operated in the data-dependent mode to simultaneously measure survey scan MS spectra (350–1,800 m/z, R = 60,000 defined at m/z 400). Up to the 20 most intense peaks were isolated and fragmented with collision-induced dissociation (CID). System controlling and data collection were carried out using Xcalibur software version 2.2 (Thermo Scientific).

### MS Data Processing

MS raw files were processed for each experiment separately with MaxQuant (version 1.5.2.8) using the integrated Andromeda search engine and UniProt FASTA database from mouse (*Mus musculus*; 2013_05). The search included variable modifications for methionine oxidation (M) and acetylation (protein N-term) as well as fixed modification for carbamidomethylation (C). Trypsin/P was set as the cleavage specificity with up to two missed cleavages allowed. The false discovery rate (FDR) cutoffs were set at 0.01 at the peptide and protein levels and the minimum peptide length was set at 7. Identification across different replicates was achieved by enabling the “match between runs” option with a matching time window of 5 min.

### Cognitive Testing

For cognitive function studies, 11.5-month-old female 3xTg-AD and NTg mice were administered vehicle (20% [w/v] 2-hydroxypropyl-β-cyclodextrin buffered with 25 mM sodium citrate pH 6.0; MilliporeSigma) or the CK1δ/ε inhibitor PF-670462 (20 mg/kg body weight/day; Cayman Chemical) s.c. daily at ZT11 in LD. Light levels during the course of the experiment were confirmed using a light sensor placed inside the housing room. Five days after the start of injections, mice underwent Y-maze and open field testing on days 6 to 9 of treatment. Mice were brought into the behavioural facility 30–60 minutes prior to testing for acclimatization to the testing environment.

#### Y-Maze

Hippocampal-dependent spatial working memory was assessed using the Y-maze spontaneous alternation behaviour test as previously described with minor modifications (Knight et al., 2014). The Y-maze apparatus consisted of three arms (38 cm long, 7.6 cm wide, and 13 cm tall) of opaque black plastic at a 120° angle from each other. Mice were placed into one of the arms of the maze and arm entries were recorded for 8 minutes as the animal freely explored all three arms. The maze was wiped clean with 70% EtOH between trials. An arm entry was defined as having all four paws in an arm. Spontaneous alternation behaviour was defined as successive entries into three different arms and expressed as a percentage of the maximum number of alternations (total number of arm entries minus 2). The total number of arm entries was also recorded as a measure of ambulatory activity and mice with fewer than six arm entries were excluded.

#### Open Field

Anxiety-like behaviour was assessed using the open field test as previously described with minor modifications (Hebda-Bauer et al., 2013). Mice were placed in the centre of an empty open field box (44 cm x 44 cm x 44 cm) made of opaque white plastic and their activity was recorded using Ethovision XT 11.5 video tracking software (Noldus Information Technology; Leesburg, VA, USA) for 5 minutes. The open field was wiped clean with 70% EtOH between trials. Time spent exploring the centre zone (24 cm x 24 cm) was recorded as a measure of anxiety level, while total distance travelled was recorded as a measure of ambulatory activity.

### Behavioural Circadian Rhythm Testing

#### Running Wheels

For behavioural circadian rhythm studies, female 3xTg-AD and NTg mice entrained to a 12-h light:12-h dark schedule from weaning were individually housed with running wheels for two weeks in LD conditions, then released into DD at 10–11 months of age. Light levels during the course of the experiment were confirmed using a light sensor placed inside each housing room. After three weeks in DD, mice were administered vehicle (20% [w/v] 2-hydroxypropyl-β-cyclodextrin buffered with 25 mM sodium citrate pH 6.0; MilliporeSigma) or PF-670462 (20 mg/kg body weight/day; Cayman Chemical) s.c. daily for 10 days at CT12, where CT was defined by the ZT of the previous LD schedule. This time point was chosen to allow all mice to receive injections at a time when it had previously been shown that PF-670462 administration has robust effects on behavioural circadian rhythms in rats (Badura et al., 2007). Running wheel activity data were collected using Wheel Manager software version 2.2 (MED Associates, Inc.; Fairfax, VT, USA). Actograms were generated and chi-square periodogram analysis performed using circadian software available online (www.circadian.org) (Chang and Guarente, 2013).

#### Dose–Response Study for PF-670462

Male 3xTg-AD mice entrained to a 12-h light:12-h dark schedule from weaning were individually housed with running wheels for two weeks in LD conditions, then released into DD at 7–8 months of age. After 16 days in DD, mice were administered vehicle (20% [w/v] 2-hydroxypropyl-β-cyclodextrin; MilliporeSigma) or PF-670462 (10 or 30 mg/kg body weight/day; Cayman Chemical) s.c. daily for 9 days at CT12, where CT was defined by the ZT of the previous LD schedule. Running wheel activity data were collected and analyzed as described above for Figure S2.

## QUANTIFICATION AND STATISTICAL ANALYSES

### Bioinformatic Analysis

Initial bioinformatic analysis was performed with Perseus (version 1.5.5.3). The raw proteomic dataset for each experiment was filtered to include only proteins quantified in at least half of samples (Q50). Hierarchical clustering analysis, using the median value of logarithmized values for the normalized light-to-heavy (L/H) ratio (for N2a samples) or label-free quantification (LFQ) intensity (for hippocampal samples) of each protein, was performed after z-score normalization of the data within Euclidean distances. Prior to principal component analysis (PCA) with MetaboAnalyst (version 4.0) (Chong et al., 2018), missing values were imputed by drawing random numbers from a normal distribution with a width parameter of 0.3 of the SD of all measured values and centre shifted towards low abundance by 1.8 times this SD. Gene Ontology (GO) and Kyoto Encyclopedia of Genes and Genomes (KEGG) pathway enrichment analyses were implemented using DAVID (version 6.8; Fisher’s exact test p ≤ 0.01 relative to the Q50 backgrounds in our datasets was considered significant) (Huang da et al., 2009). Protein–protein interaction (PPI) networks were created using the STRING database (Szklarczyk et al., 2015) (confidence score cutoff = 90%) and visualized with NetworkAnalyst (Xia et al., 2015).

### Statistical Analysis

*In vitro* CK1δ/ε inhibition proteomic data were analyzed using unpaired two-tailed Student’s *t*-test (permutation-based FDR = 0.05; s0 = 0.1) for differential expression analysis. *In vivo* CK1δ/ε inhibition proteomic data were analyzed using unpaired two-tailed Student’s *t*-test (α = 0.05; log_10_ fold change [experimental/control] greater than log_10_1.2 or less than –log_10_1.2) for differential expression analysis. Y-maze and open field data were analyzed using unpaired two-tailed Student’s *t*-test. Running wheel data were analyzed by chi-square periodogram analysis (α = 0.05), and one-way analysis of variance (ANOVA) with Fisher’s least significant difference (LSD) *post hoc* test, or two-tailed unpaired or paired Student’s *t*-tests (with Welch’s correction where indicated), were performed to compare group differences as appropriate. Statistical analyses were carried out using Perseus (version 1.5.5.3) for proteomic data or Prism 6 (GraphPad; La Jolla, CA, USA) for behavioural data. Data in bar graphs are represented as means + SEM.

## DATA AND SOFTWARE AVAILABILITY

The mass spectrometry proteomics data have been deposited to the ProteomeXchange Consortium via the PRIDE partner repository with the dataset identifiers PXD012281 and PXD012282.

