## Supplemental Information for "Therapeutic Targeting of Casein Kinase 1δ/ε in an Alzheimer’s Disease Mouse Model"

Supplemental Information includes two figures and six tables.

Figure S1. PF-670462 Administration Alters Expression of Proteins Involved in Long-Term Potentiation and Amyloid Precursor Protein Processing in the Hippocampus of 3xTg-AD Mice, Related to Figures 2 and 3.

Figure S2. Dose Determination for PF-670462 Running Wheel Studies in 3xTg-AD Mice, Related to Figure 5.

Table S1. List of Proteins Accurately Quantified in Neuro-2a Cells, Related to Figure 1.

Table S2. List of Proteins Differentially Expressed with PF-670462 Treatment in Neuro-2a Cells, Related to Figure 1.

Table S3. List of Hippocampal Proteins Accurately Quantified for CT10, Related to Figure 2.

Table S4. List of Hippocampal Proteins Accurately Quantified for CT14, Related to Figure 2.

Table S5. List of Proteins Differentially Expressed with PF-670462 Administration in the Hippocampus of 3xTg-AD Mice for CT10, Related to Figures 2 and 3.

Table S6. List of Proteins Differentially Expressed with PF-670462 Administration in the Hippocampus of 3xTg-AD Mice for CT14, Related to Figures 2 and 3.

### SUPPLEMENTARY FIGURES

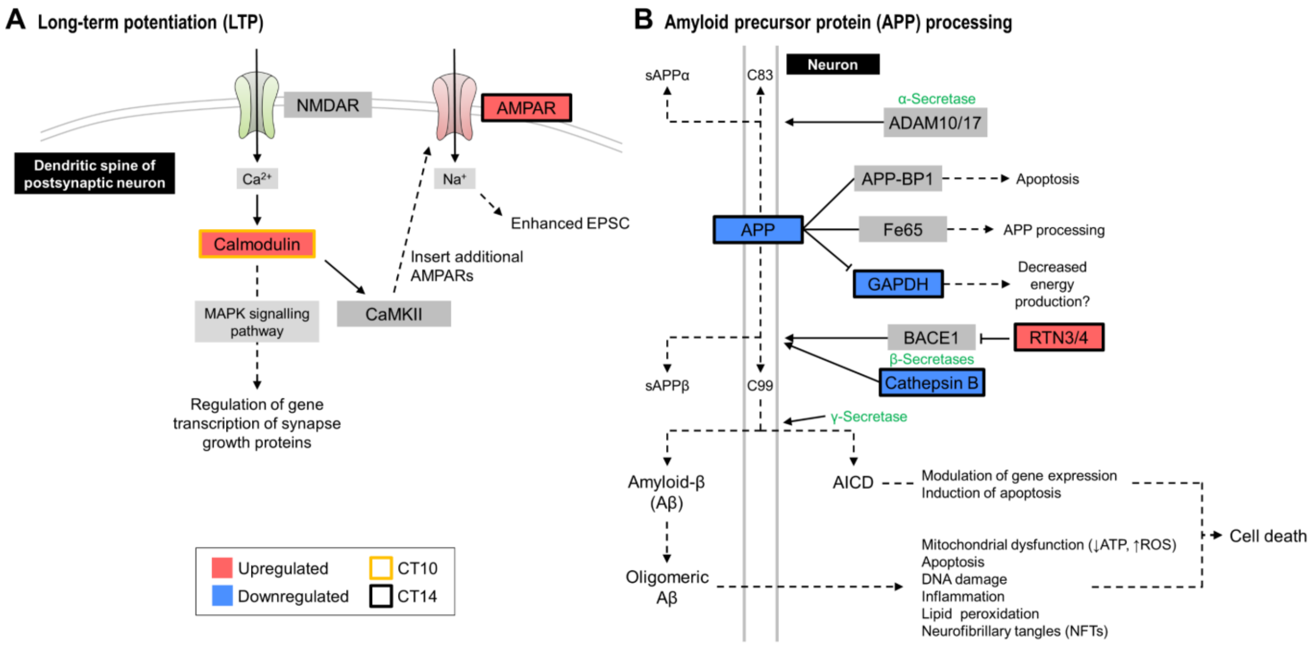


**Figure S1. PF-670462 Administration Alters Expression of Proteins Involved in Long-Term Potentiation and Amyloid Precursor Protein Processing in the Hippocampus of 3xTg-AD Mice, Related to Figures 2 and 3**

8-month-old NTg and 3xTg-AD mice were treated daily with PF-670462 (30 mg/kg/d) or vehicle for 20 days, then hippocampal tissues were collected at two time points for proteomic analysis. Protein extracts were digested with trypsin and analyzed by LC-MS/MS. For CT10: n = 3 NTg vehicle; n = 4 3xTg-AD vehicle; n = 3 3xTg-AD PF-670462. For CT14: n = 4 NTg vehicle; n = 5 3xTg-AD vehicle; n = 4 3xTg-AD PF-670462.

(A and B) Schematic diagrams depicting the A) hippocampal LTP and B) APP processing pathways, highlighting proteins that were differentially expressed with PF-670462 administration ($p < 0.05$, two-tailed unpaired Student's *t*-test; $\log_{10}$ fold change > $\log_{10}1.2$ or < $-\log_{10}1.2$) Proteins depicted in gray were either not accurately quantified in our datasets or not differentially expressed with PF-670462. Adapted from the Kyoto Encyclopedia of Genes and Genomes (KEGG) pathways 04720 (long-term potentiation) and 05010 (Alzheimer disease). ADAM, disintegrin and metalloproteinase domain-containing protein; AICD, APP intracellular domain; AMPAR, α-amino-3-hydroxy-5-methyl-4-isoxazolepropionic acid receptor; APP, amyloid precursor protein; APP-BP1, APP binding protein 1; BACE1, β-site APP cleaving enzyme; CaMKII, $Ca^{2+}$/calmodulin-dependent protein kinase II; EPSC, excitatory postsynaptic current; Fe65, APP binding family B member 1; GAPDH, glyceraldehyde-3-phosphate dehydrogenase; MAPK, mitogen-activated protein kinase; NMDAR, *N*-methyl-D-aspartate receptor; ROS, reactive oxygen species; RTN, reticulon.

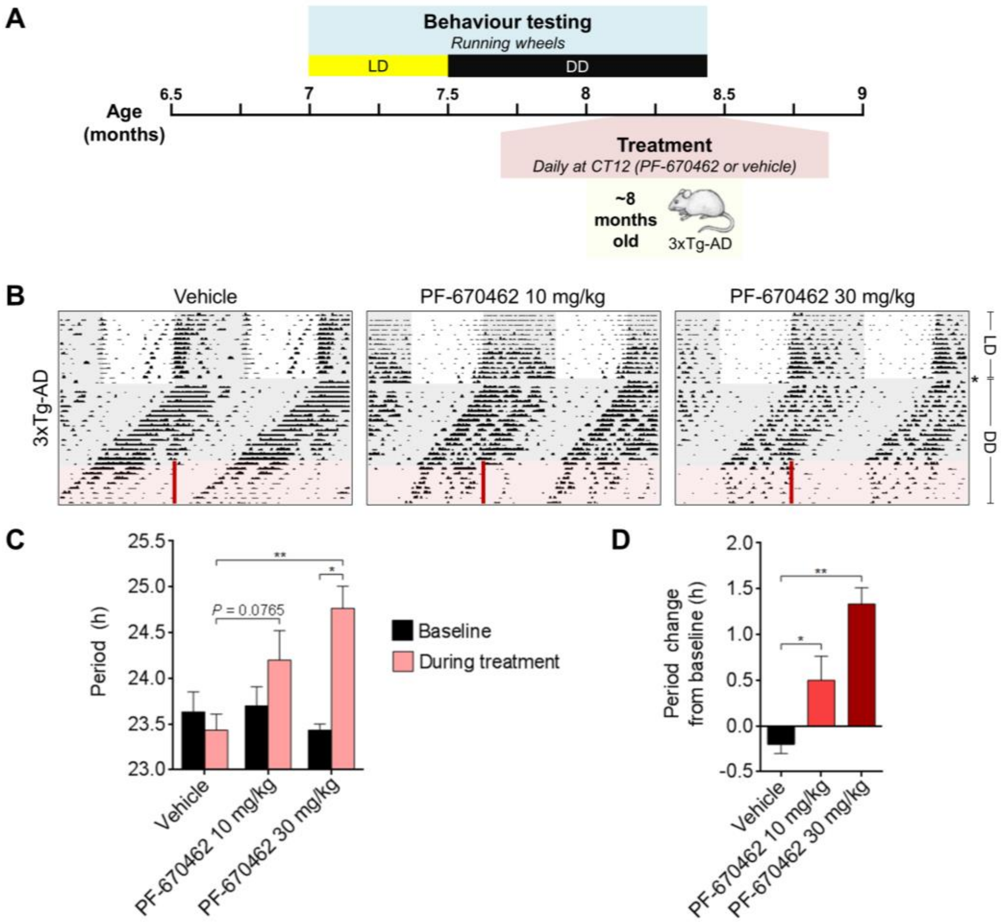


**Figure S2. Dose Determination for PF-670462 Running Wheel Studies in 3xTg-AD Mice, Related to Figure 5**

(A) Study design: Wheel-running activities of 3xTg-AD mice were recorded for 2 weeks in LD, then mice were released into DD at ~7.5 months of age. After 16 days in DD, mice received daily s.c. injections of PF-670462 (10 or 30 mg/kg/d) or vehicle for 9 days in DD. n = 3 mice per treatment group.

(B) Representative actograms of wheel-running activities under different lighting conditions of ~8-month-old 3xTg-AD mice treated s.c. daily with vehicle, PF-670462 10 mg/kg/d, or PF-670462 30 mg/kg/d. The timing of treatment is indicated by red bars. Missing data due to computer failure are indicated by an asterisk.

(C) Free-running period under DD before and during treatment of 3xTg-AD mice with vehicle, PF-670462 10 mg/kg/d, or PF-670462 30 mg/kg/d (**$p < 0.01$ versus vehicle during treatment, one-way ANOVA with Fisher's LSD *post hoc* test; *$p < 0.05$ versus baseline, two-tailed paired Student's *t*-test).

(D) Change in free-running period under DD from pre-treatment baseline (*$p < 0.05$, **$p < 0.01$ versus vehicle, one-way ANOVA with Fisher's LSD).

Data are represented as means + SEM.

### SUPPLEMENTAL TABLE TITLES AND LEGENDS

**Table S1. List of Proteins Accurately Quantified in Neuro-2a Cells, Related to Figure 1.** All proteins in this list were quantified in at least half of samples (Q50). The majority protein IDs, protein names, gene names, Gene Ontology biological process (GOBP) functional annotations, and logarithmized light-to-heavy (L/H) ratios are supplied.

**Table S2. List of Proteins Differentially Expressed with PF-670462 Treatment in Neuro-2a Cells, Related to Figure 1.** All proteins in this list were quantified in at least half of samples (Q50) and differentially expressed in Neuro-2a cells treated for 24 hours with PF-670462 (5 µM) versus DMSO (FDR-corrected $p < 0.05$, two-tailed unpaired Student's *t*-test; $s0 = 0.1$). The majority protein IDs, protein names, gene names, Gene Ontology biological process (GOBP) functional annotations, and logarithmized light-to-heavy (L/H) ratios are supplied.

**Table S3. List of Hippocampal Proteins Accurately Quantified for CT10, Related to Figure 2.** All proteins in this list were quantified in at least half of samples (Q50). The majority protein IDs, protein names, gene names, Gene Ontology biological process (GOBP) functional annotations, and logarithmized label-free quantification (LFQ) intensities are supplied.

**Table S4. List of Hippocampal Proteins Accurately Quantified for CT14, Related to Figure 2.** All proteins in this list were quantified in at least half of samples (Q50). The majority protein IDs, protein names, gene names, Gene Ontology biological process (GOBP) functional annotations, and logarithmized label-free quantification (LFQ) intensities are supplied.

**Table S5. List of Proteins Differentially Expressed with PF-670462 Administration in the Hippocampus of 3xTg-AD Mice for CT10, Related to Figures 2 and 3.** All proteins in this list were quantified in at least half of samples (Q50) and differentially expressed in the hippocampus of 3xTg-AD mice treated for 20 days with PF-670462 (30 mg/kg/d) versus vehicle ($p < 0.05$, two-tailed unpaired Student's *t*-test; $\log_{10}$ fold change $> \log_{10}1.2$ or $< -\log_{10}1.2$). The majority protein IDs, protein names, gene names, Gene Ontology biological process (GOBP) functional annotations, and logarithmized label-free quantification (LFQ) intensities are supplied.

**Table S6. List of Proteins Differentially Expressed with PF-670462 Administration in the Hippocampus of 3xTg-AD Mice for CT14, Related to Figures 2 and 3.** All proteins in this list were quantified in at least half of samples (Q50) and differentially expressed in the hippocampus of 3xTg-AD mice treated for 20 days with PF-670462 (30 mg/kg/d) versus vehicle ($p < 0.05$, two-tailed unpaired Student's *t*-test; $\log_{10}$ fold change $> \log_{10}1.2$ or $< -\log_{10}1.2$). The majority protein IDs, protein names, gene names, Gene Ontology biological process (GOBP) functional annotations, and logarithmized label-free quantification (LFQ) intensities are supplied.
